# Short communication: Oral microbiome as a potential proxy for grazing livestock methane emissions

**DOI:** 10.64898/2026.03.26.714587

**Authors:** Chian Teng Ong, Tony Cavallaro, Yuhao Li, Alana C Boulton, Basil S Firewski, Marloes D Nitert, Kieren D McCosker, Sam Clark, Scott Cullen, Melissah Dayman, Milou H Dekker, Paul Gangemi, Kerry Goodwin, Tim Grant, Rachelle Hergenhan, David Johnston, Natalie Scott, Bradley Taylor, Cameron Whistler, Ben J Hayes, Marina R S Fortes, Elizabeth Ross

## Abstract

Enteric methane emissions from ruminant livestock contribute to global warming, creating an urgent need for effective mitigation strategies that do not compromise animal productivity and welfare. Methanogenic archaea within the rumen microbiome drive enteric methane emissions. However, large-scale rumen-fluid sampling in commercial production systems is impractical, due to its invasive nature and the associated logistical challenges. This study hypothesised that rumination enables the capture of rumen microbial signals within the oral cavity and using oral microbiome profiles to provide a practical, non-invasive alternative method for proxy methane phenotyping in commercial production systems. To test the hypothesis, we estimated the oral microbiability, defined as the proportion of phenotypic variance in methane emissions explained by oral microbiome variation. Samples were collected from 209 animals across two trials in Queensland, Australia. Oral microbiome samples were obtained from all animals, with paired rumen samples in one trial, and methane emissions were measured using either the sulphur hexafluoride (SF_6_) tracer technique or the GreenFeed system. Microbial features were characterised using taxonomic and functional annotations, and microbiability was estimated using mixed linear models incorporating microbiome-based relationship matrices. Although the small sample size limited strong conclusions, the oral microbiability estimates reported in this study were comparable to those derived from rumen samples. Functional microbial profiles generally explained a greater proportion of methane variation than taxonomic profiles, suggesting that microbial function is more closely linked to methane production than community composition alone. However, these differences were not statistically significant due to large standard errors. These findings suggest that oral microbiome sampling potentially provides a practical, minimally invasive, scalable proxy method for methane emissions of individual cattle in grazing systems, where direct methane gas measurements are labour-intensive and difficult to implement. Integrating oral microbiome profiles in the existing breeding model with the host genetics, weight and environmental factors could provide a promising pathway for enabling selection for low emissions and advancing reduced emissions livestock farming under real-world production conditions.

**Lay summary:** Cattle produce methane as part of their normal digestion and this contributes to climate change. Reducing methane emission in grazing livestock systems is therefore important. However, measuring methane from individual grazing animals is difficult, costly, and often impractical under commercial conditions. The rumen microbiome has been used as a proxy for estimating methane emissions, but collecting rumen samples is invasive and impractical for large-scale use. Because rumination transfers material from the rumen to the mouth, we investigated whether microbes found in cattle mouths could also be used to estimate how much methane an individual animal produced. We suggest that mouth-swab sampling method can be an alternative to rumen fluid sampling because it was less invasive, relatively quick and practically applicable in commercial conditions. Importantly, the microbiome explained a meaningful proportion of the between-animal variation for methane emission. This suggests that collection of mouth swabs is a potentially scalable alternative proxy method to identify cattle that naturally produce less methane. Overall, our findings support the potential use of oral ruminant microbial information to improve breeding and management strategies aimed at reducing methane emissions while maintaining productive livestock systems.

**Teaser Text:** This study demonstrates that collecting oral swabs from the mouths of grazing beef cattle could provide a scalable method to estimate individual methane emissions in commercial production systems, offering a practical alternative to invasive rumen sampling and complex gas measurement systems. These findings support the development of scalable breeding and management strategies for methane mitigation in large-scale livestock production systems.

## Introduction

Methane is a strong greenhouse gas (Moss et al., 2000), and ruminant enteric fermentation contributed to ∼30% of global methane emissions (FAO, 2023). Research efforts to reduce methane from livestock have increasingly focused on breeding ruminant livestock with lower methane emissions (Manzanilla-Pech et al., 2021; Saborío-Montero et al., 2021; Manzanilla-Pech et al., 2022). However, accurately measuring and tracking enteric methane emissions at the individual animal level in large enough numbers to inform breeding programs remains a significant challenge. Technologies, such as respiration chambers, the GreenFeed system and the SF_6_ tracer gas methodology, are employed to quantify methane emissions, but they are either low throughput, labour-intensive, costly or impractical for use on an industry-wide scale (Garnsworthy et al., 2019). This limits the development and the precision of methane mitigation strategies, including development of estimated breeding values to select for low-emission livestock as well as large scale monitoring programs to validate the impact of methane reductions technologies.

The rumen microbiome has been extensively investigated because it harbours methanogenic archaea, the primary biological agents of enteric methane emissions, which exist in mutualistic symbiosis within the rumen. The methanogenic archaea facilitate the breakdown of indigestible fibre and supporting the dietary and metabolic requirements of the ruminant host (Bergman, 1990). Methods that use the entire microbiome, often fitted as a relationship or correlation matrix, have demonstrated a statistically significant association between methane yield predicted from the rumen microbiome and methane yield measured using classic respiration chambers or portable accumulation chambers (Ross et al., 2013; Hess et al., 2023; Li et al., 2025). Hence, sampling the rumen microbiome has been proposed as an alternative or proxy to measuring methane gas directly.

Microbiability, which represents the proportion of variation in methane-related traits explained by differences in the microbiome, has been approximated at 13% to 40% when using the rumen microbiome as a predictor of methane production (Difford et al., 2018; Li et al., 2019). Positive rumen microbiability indicates the predictive potential of the rumen microbiome to estimate methane production (González-Recio et al., 2023; Sepulveda et al., 2025; Waters et al., 2025). Nonetheless, collecting rumen fluid poses a substantial challenge to fully leveraging the potential of rumen microbiome data. Common rumen fluid sampling methods, such as ruminal fistulation, esophageal tubing, or rumenocentesis, are invasive, slow to implement, pose risks to animal health and require skilled technicians to implement (Nordlund and Garrett, 1994; Muizelaar et al., 2020; Castillo and Hernández, 2021). For these reasons, these methods are impractical for large-scale deployment in commercial farm settings. Therefore, the rumen microbiome alone is unsuitable for microbiome-based methane predictions where large datasets are required to account for multiple sources of variation, including breed, climate, geographical region, feed type and forage quality, especially in non-research herds.

Building on the initial work using rumen microbiome to predict enteric methane production, oral samples were proposed as a more user-friendly and less invasive sampling method, based on the assumption that rumen microbiota are deposited in the mouth during rumination (Ross, 2013). Multiple studies have verified the use of oral samples as a practical, minimally invasive and scalable alternative to rumen fluid sampling because oral samples yield a similar microbiome profile to rumen samples, despite fewer archaea detected in oral samples (Tapio et al., 2016; Marcos et al., 2024). Additionally, there are promising examples of using oral microbiome as a proxy to rumen data for predicting feed intake (Marcos et al., 2024) and as a proxy for subacute rumen acidosis (Liu et al., 2025). In this study, we examine the proportion of between-animal variation for methane production that can be attributed to oral microbiome variation, or oral microbiability, of cattle under grazing conditions.

## Methods and Materials

### Ethics declarations

All procedures involving animal use were approved under Animal Ethics Approval AE000438 and AE000657 by the Animal Ethics Committee of the University of Queensland (UQ). All protocols in this study were performed in accordance with the UQ Animal Ethics Committee approved standard operating procedures (SOPs) for Companion and Production Animals. The study was carried out in compliance with the ARRIVE guidelines.

### Animal recruitment

A total of 209 animals were enrolled across two trials. The first trial, “QASP” trial, was conducted at QASP (Queensland Animal Science Precinct) at the UQ Gatton Campus (n = 38). The second trial, “Extensive” trial, was an extensive grazing trial comprising cattle from three grazing stations in Queensland, Australia: Brian Pastures Research Facility (n = 19), Spyglass Beef Research Facility (n = 76), and Goldsborough Station (n = 95).

For the “QASP” trial, two biological sample types—oral and rumen—were collected for microbiome characterisation. Sampling was conducted across four seasons (Winter 2023, Spring 2023, and Summer 2024, Autumn 2024), yielding longitudinal samples from the same cohort of 38 grazing Brahman heifers at four time points. All animals in the “QASP” trial were monitored for enteric methane emissions using the sulfur hexafluoride (SF_6_) tracer technique. For animals enrolled in the “Extensive” trial, only oral samples were collected for microbiome analysis, and methane emissions data were obtained using the GreenFeed system. Further details of both trials are provided below.

### Oral sample collection

Oral samples were collected using an in-house swab collection prototype (Figure 1). Briefly, the animal was guided into a veterinary squeeze chute. Following weigh measurement, the head of the animal was then restrained using the bail in an elevated position. An oral sample was then collected using the oral swab collection prototype, which was inserted into the oral cavity and brushed against the buccal surface, sublingual area and palate for approximately 60 seconds in total. The collected swab was snipped and kept in the collection tube with 2ml of sterile storage buffer made of phosphate-buffered saline and 30% glycerol. The collected samples were kept on ice during transportation and -80°C for long-term storage prior to microbiome DNA extraction.

**Figure 1.**
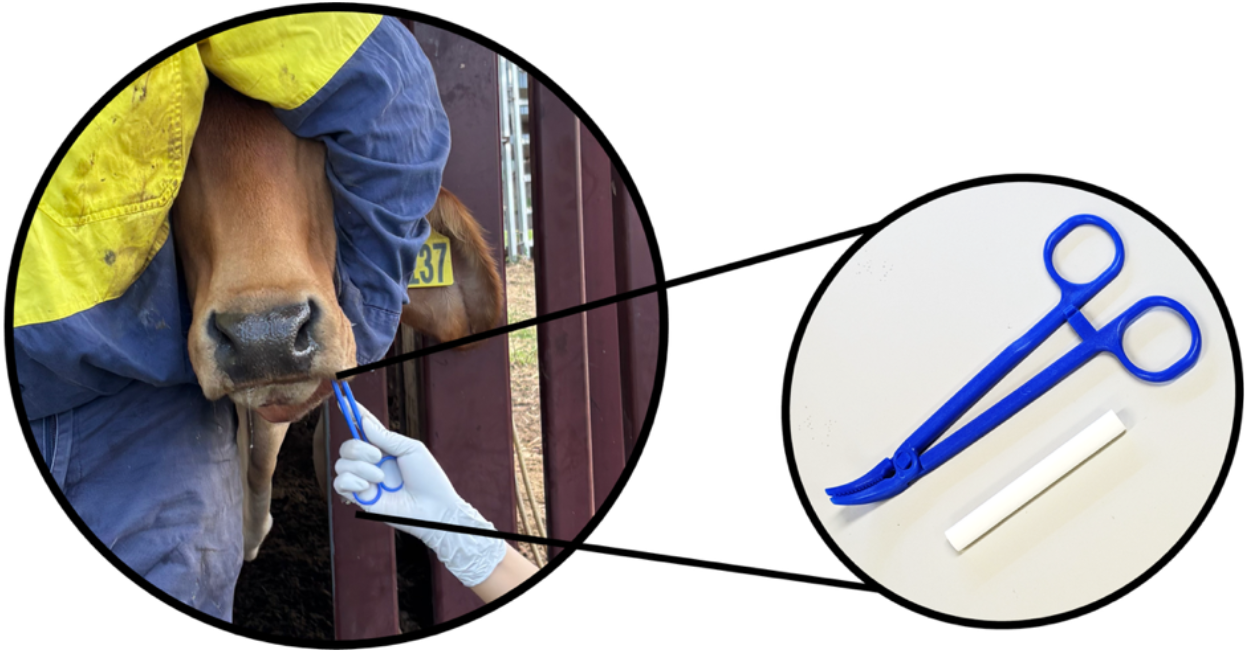
In-house oral swabbing tool comprising disposable plastic pliers and a cotton swab.

### Rumen sample collection

Rumen samples were collected after oral sampling to minimise cross-contamination. To obtain rumen contents, a T-bar metal tube was inserted into the rumen by an experienced technician of the QASP team. A flexible hose was then connected between the tube and a mechanical pump. Once the equipment was in place and the animal was settled, rumen contents were pumped out. The collected material was filtered through a metal sieve and transferred into 50 mL Falcon tubes. Samples were kept on ice during transport and subsequently stored at −80 °C for long-term preservation.

### Sulphur hexafluoride (SF_6_) methane phenotype measurement

In the “QASP” trial, a protocol for measuring enteric methane emission in grazing cattle using SF_6_ gas methodology was used and is described in detail by (Boulton et al., 2025). Briefly, the enteric methane of 38 commercial Brahman heifers (18–24 months) was quantified using the SF_6_ tracer gas technique at QASP research facility. Permeation tubes were manufactured and calibrated according to established guidelines prior to oral administration (Boulton et al., 2025). Breath samples were collected using fitted saddles and halters, which were adapted from original designs to include coiled recoil tubing, thereby facilitating grazing movement and reducing equipment failure (Boulton et al., 2025). The study targeted five consecutive 24-hour collection days per sampling period across different seasons. Background methane and carbon dioxide levels were corrected using four spatially distributed environmental canisters, with sample aliquoting performed in open-air cattle yards to eliminate laboratory contamination with SF_6_ gas before gas chromatography analysis. The entire methodology was described in detailed by Boulton et al. (2025).

### Greenfeed methane phenotype measurement

In the “Extensive” trial, methane (CH_4_) and carbon dioxide (CO_2_) emissions were measured using GreenFeed units under representative grazing systems and the trial is described in detail by (Whistler et al., 2026). Briefly, animals underwent a three-week acclimation period prior to data collection to familiarise them with the units, during which lucerne pellets were provided as an attractant and dispensed at an increased frequency to encourage correct head positioning for breath sampling. To promote uniform acclimation, animals that repeatedly used the units were temporarily removed and reintroduced after five days to allow less responsive animals to adapt. During the trial, cattle had *ad libitum* access to forage and GreenFeed units, with sampling permitted once every two hours and pellet delivery limited to a maximum of eight visits per animal per day. Only visits exceeding two minutes of continuous recording were retained for analysis, and no supplementary feeding was provided beyond the pellets dispensed by the units.

### Microbiome DNA extraction

The oral swab samples were thawed and transferred into a 10 mL syringe barrel, and the material was expressed by compressing the plunger to recover the collected oral fluids. For rumen samples, 10 mL of material was taken directly from the storage tube after gentle mixing and used for DNA extraction.

To extract the total DNA, the samples were first centrifuged at 4,000 *g* at 4 °C for 15 minutes, and the supernatant discarded. The pellets were extracted for cattle oral microbiome DNA using the QIAamp PowerFecal Pro DNA Kit (QIAGEN, Germany) following the manufacturer’s instruction with an additional proteinase K step. Briefly, the pellets were suspended in the lysis buffer and proteinase K before mechanical lysis with bead beating. Then the samples were incubated at 56 °C for 60 minutes, followed by DNA precipitation, washing and elution as outlined in the manufacturer’s protocol. The quantity and the quality of the DNA were assessed using using the Qubit™ 4 Fluorometer (Thermo Scientific, USA) and NanoDrop 1000 Spectrophotometer (Thermo Scientific, USA).

### Microbiome DNA sequencing and basecalling

Barcoded DNA libraries were prepared using the Native Barcoding Kit 96 V14 SQK-NBD114.24 (Oxford Nanopore Technology, UK) from the extracted microbiome DNA with a minor change on the incubation time. Briefly, 400ng DNA of each sample were added to the end-repair reaction mix and incubated for an extended duration of 20 minutes prior to barcode ligation. The barcoded DNA samples were subsequently pooled for adaptor ligation, after which the adaptor-ligated pooled DNA libraries loaded onto a PromethION flow cell FLO-PRO114M with R10.4.1 chemistry (Oxford Nanopore Technology, UK) prior to sequencing with PromethION 2 Solo (Oxford Nanopore Technology, UK). Each sample was sequenced to a depth of at least one million reads. Where a sample that did not reach this threshold due to barcode imbalance the samples were re-sequenced until there was at least 1 million reads for each sample. All 342 oral and 152 rumen samples achieved the targeted 1 million reads. The raw data was transferred to the Linux system for super accurate (SUP) model basecalling using Dorado v0.6.2 (Oxford Nanopore Technology, UK).

### Quality filtering and metagenomic analyses

Since oral samples are often saturated with host genetic materials, the bovine reads in the oral samples of this study were set aside by mapping the base-called reads against two bovine genomes, including ARS-UCD1.2 *Bos taurus* genome (GCA_002263795.2) and Brahman genome (Ross et al., 2022), using Minimap2 2.17 (r941) (Li, 2018). Reads which were not mapped to the bovine genomes were selected for quality filtering using Chopper v0.10.0b (De Coster and Rademakers, 2023) to remove the first 15 bases of every read as well as reads with less than 7 in quality score and less than 500 bases. The dataset of each sample were examined using NanoPlot 1.3.0 (De Coster et al., 2018) at every step.

The metagenomic analyses were conducted on a fully automatic pipeline, SqueezeMeta v1.6.5 (Tamames and Puente-Sánchez, 2019), which is equipped with several databases including GenBank nr (Benson et al., 2016) for taxonomic assignment, and the Kyoto Encyclopedia of Genes and Genomes (KEGG) database (Kanehisa and Goto, 2000) and the Clusters of Orthologous Groups (COG) database (Tatusov et al., 2000) via eggNOG 5.0 (Huerta-Cepas et al., 2019) for functional assignment. The alternative script sqm_longreads.pl was applied to perform the metagenomic classification using Diamond Blastx (Buchfink et al., 2014) on the filtered reads in each sample dataset. Subsequently, the combine-sqm-tables.py script was used to combine the individual results into one dataset for downstream analyses in R studio (RStudio Team, 2020) using R packages phyloseq (McMurdie and Holmes, 2013), vegan (Jari Oksanen et al., 2020) and ggplot2 (Wickham., 2016). For taxonomic assessment, OTU which were assigned as “unclassified”, “uncultured” and “unknown” as taxonomic levels up to Family were filtered out.

### Microbiability

The microbiome abundance matrix was constructed by extracting, for each sample, the sample identifier, the corresponding taxonomic or functional annotation, and the total abundance associated with each assignment. The abundance matrix was transformed first by adding a pseudo count of 1 to the total abundances of all the OTU entries to avoid zeros during downstream transformation, and second by centred log-ratio (CLR) transformation to account for the compositional differences of the samples. A microbial relationship matrix *G*_m_ was constructed based on the transformed abundance matrix, M, using rrBLUP package (Endelman, 2011) as:

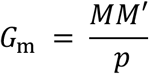

The proportion of variance in methane intensity explained by the oral microbiome (microbiability) was calculated using ASReml-R (Butler et al., 2017) using the following mixed model:

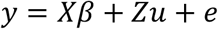

In the “QASP” trial, *y* represents the mean of valid daily methane emission measurements (g CH_4_), measured using the SF_6_ technique over a five-day period. As each animal in this trial was sampled across four seasonal time points, an animal-level microbiome relationship matrix was constructed by aggregating the sample-level microbiome relationship matrix. This was done by averaging all pairwise sample similarities within and between animals, producing a single microbiome relationship value for every animal pair. For this dataset, the fixed effect vector, *β*,included season, pregnancy status and liveweight. Liveweight was fitted as a fixed effect to account for variation in methane emissions attributable to differences in animal size and associated metabolic demand. The random effect was modelled as 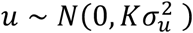 where K is the animal-level microbiome relationship matrix.

For the “Extensive” trial, in which each animal contributed only a single microbiome sample, the microbiome relationship matrix was used at the sample level. For this dataset, the fixed-effect vector β included site and liveweight. “Site” was included as a fixed effect to account for systematic environmental or management differences between the different sampling sites while liveweight is fitted in the model to account for the methane variations attributed to body size and associated metabolic demand. The random animal microbiome effect was modelled as 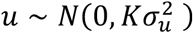, where K is the sample-level microbiome relationship matrix.

In both datasets, the residual errors were 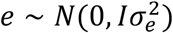. Variance components 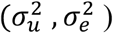 were extracted from the fitted model. Microbiability was calculated as:

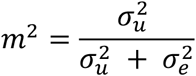

## Results

The sequencing output of the bovine oral samples collected from cattle under the “QASP” trial averaged at 1,851 Mb per sample, while the rumen samples yielded an average of 3,063.40 Mb per sample (Supplementary Table 1). After quality filtering and host-read removal, the retained non-host microbial reads were reduced to 338.29 Mb for the oral samples and 1,497.82 Mb for the rumen samples. For the oral samples collected from the cattle stations, the sequencing output averaged 2,138.20 Mb per sample, with 226.33 Mb retained after quality filtering and removal of host-derived reads. Across all oral samples in this study, approximately 80–90% of total reads were identified as cattle-derived and removed during host read filtration.

For the “QASP” trial, the microbiability estimates were obtained for methane phenotypes measured using the SF_6_ tracer technique, based on rumen and oral microbiome profiles at different functional and taxonomic levels. At the genus (taxonomic) level, the oral microbiome explained a greater proportion of phenotypic variance compared with the rumen microbiome (Table 1). This pattern was consistent also for functional annotations. Using COG functional assignment, the oral samples showed higher microbiability than rumen samples. Similarly, KEGG functional estimates demonstrated that oral microbiome features explained more methane variation than the rumen microbiome. There was no significant difference between annotation methods; however, across trials, oral samples tended to have higher microbiability estimates, though the relatively large standard deviations indicate that these differences were not statistically robust, suggesting comparability with rumen samples.

**Table 1.**
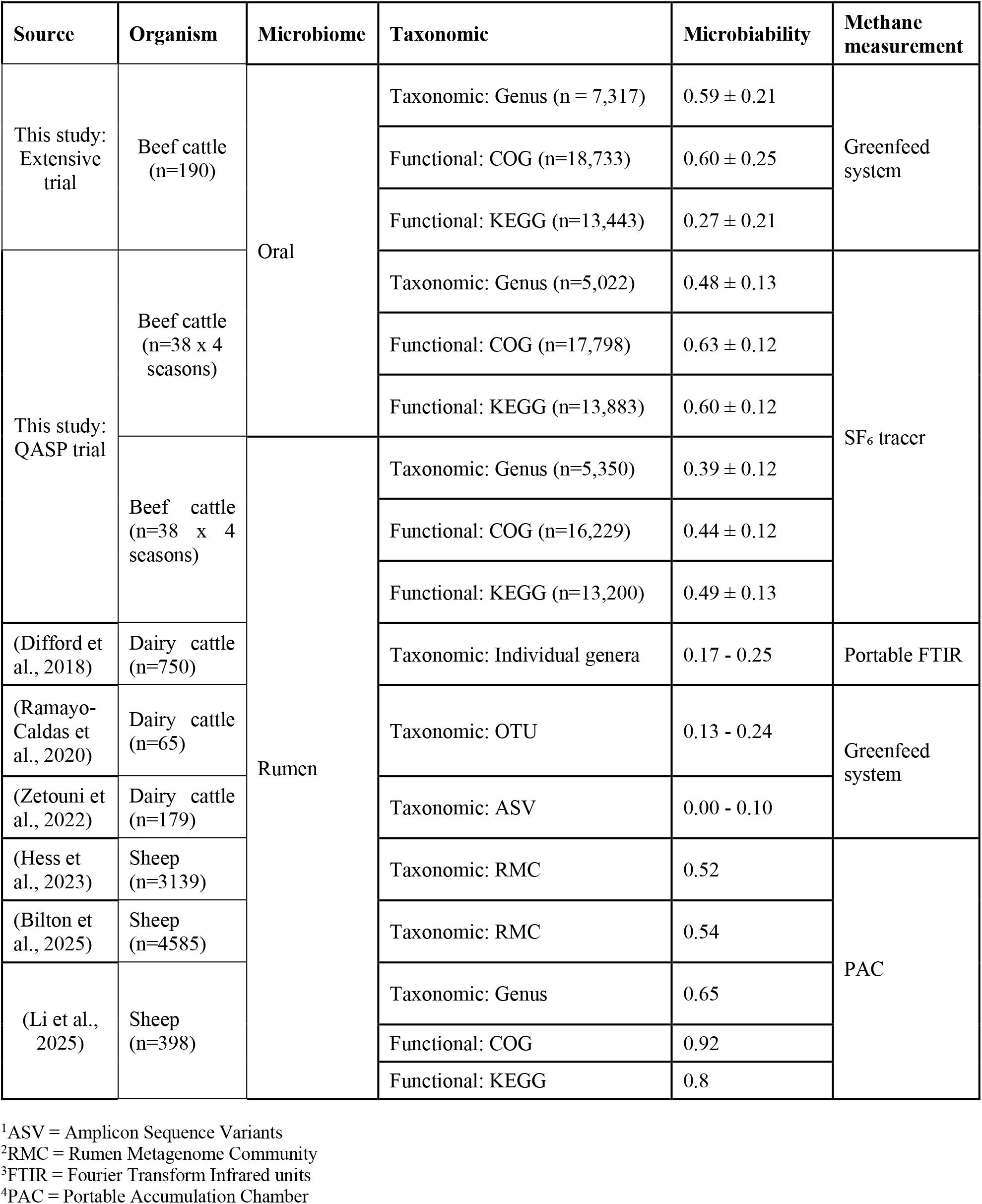
Microbiability (m^2^) estimates for methane emissions based on taxonomic and functional microbial signatures from rumen and oral samples, using methane phenotypes measured by different methane measurement technologies reported in this study and other published studies. Microbiability estimates from the current study are presented as estimate ± standard error.

In the “Extensive” trial, microbiability estimates based on genus-level (taxonomic) microbiome profiles were high; however, the associated standard errors were larger than those observed in the QASP trial (Table 1). A similar pattern was observed for COG functional annotations, where microbiability remained high. In contrast, KEGG-level features showed a lower microbiability, indicating limited precision and weaker support for a strong microbial contribution at this functional level. Nonetheless, the relatively high standard errors across all microbiome classification type in the “Extensive” trial reflect the high variability in the microbial signal strength and a considerable uncertainty around the estimate.

## Discussion

In this study, we demonstrated the proportion of enteric methane emissions explained by the oral and rumen microbiome structure (microbiability) in cattle across two trials, namely the “QASP” trial and the “Extensive” trial. We observed microbiability estimates ranging between 0.27 and 0.63 in this study. This suggests that oral microbiome profiles have the potential to be used as a proxy for methane emissions, specifically, the component of methane emissions that is independent of liveweight, given that liveweight was fitted as a covariate in the models. Under these conditions, prediction accuracies as high as 0.7 may be achievable given that the square root of microbiability represents the upper bound of prediction accuracy (Ould Estaghvirou et al., 2013; Ross and Hayes, 2022). Nevertheless, this estimate is derived indirectly rather than from empirical prediction performance, as achieving such accuracy would likely require a substantially larger reference population than that used in the present study.

In the “QASP” trial, where both rumen and oral samples were available to estimate microbiability using the same phenotype set measured with the SF_6_ tracer gas methodology, there was no disadvantage to using oral samples over rumen samples to capture methane phenotype variation. Notably, the oral microbiability estimates reported here were higher than previously reported rumen microbiability estimates in dairy cattle where methane was measured using GreenFeed units (Ramayo-Caldas et al., 2020; Zetouni et al., 2022) and portable Fourier transform infrared units (Difford et al., 2018) (Table 1). In addition, the oral microbiability observed in beef cattle was comparable to rumen microbiability estimates reported in sheep measured using portable accumulation chambers (Hess et al., 2023; Bilton et al., 2025; Li et al., 2025). In the “Extensive” trial, where we estimated the oral-based microbiability using the GreenFeed systems under extensive grazing conditions, the oral microbiome was also capable of explaining approximately 60% of the methane yield variation. Taken together these results suggest that there is merit in further studies which directly calculate the predictive performance of the oral microbiome for enteric methane emissions.

### Measurement error could affect microbiability

Differences in microbiability estimates between methane measurement methods highlight the sensitivity of microbiome–phenotype associations to phenotype definition, measurement precision, and statistical uncertainty. Disagreement among methane measurement methods has been widely documented (Hristov et al., 2016; Doreau et al., 2018; Rey et al., 2019; Wallace et al., 2019; Ma et al., 2024). Importantly, the magnitude and direction of these differences are often inconsistent over time, largely due to the inherent low repeatability of the complex methane measurement approaches (Hristov et al., 2016). Both the SF_6_ tracer gas methodology and the GreenFeed system have been classified as having only moderate repeatability by experienced experts (Garnsworthy et al., 2019). Microbiability estimates derived from SF_6_ tracer measurements were consistently moderate to high across sample types and microbial classifications, whereas GreenFeed-based estimates were more variable, although differences were not statistically significant due to large standard errors. This variability likely reflects methodological differences between the two systems. The SF_6_ methodology provides continuous, integrated measurements of methane emissions over 5 days (Boulton et al., 2025) which likely leads to more precise phenotype measurement, while GreenFeed systems rely on short-duration spot measurements and requires longer measuring periods (up to 45 days) to achieve acceptable repeatability as the measurements are more susceptible to animal behaviour and visit frequency (Arbre et al., 2016; Ryan et al., 2022). Future studies should examine the microbiability of the cattle oral microbiome in relation to methane emission phenotypes measured using different technologies to enable robust comparison. This is especially relevant in the context of economic trade-offs with accuracy for phenotyping methods used to build prediction equations.

### Oral samples as a potential proxy for rumen samples

Interestingly, in the “QASP” trial the oral microbiability was similar to the rumen microbiability. The rumen microbiome has traditionally been regarded as the primary biological indicator of enteric methane emissions, with oral sampling initially proposed as a practical alternative based on cattle rumination behaviour. Accordingly, oral microbiability was hypothesized to be similar to or lower than that of rumen-based estimates. However, we did not observe this disadvantage. This adds weight to the idea that oral sampling could be a welfare friendly alternative to rumen sampling for methane prediction, especially in an applied setting. Notably, associations between methane emissions and milk or faecal-based biomarkers have also been reported (Chilliard et al., 2009; Dijkstra et al., 2011; Manafiazar et al., 2021; Youngmark and Kraft, 2025). However, milk is not a feasible method for grazing beef cattle, and the physical oral-rumen link could provide a more representative proxy for rumen samples compared to faeces which are separated by the gastric stomach and therefore do not have a direct connection to the rumen. The caveat to this idea is that oral samples still require significant animal handling and restraint in a crush, whereas faecal samples can be passively obtained without the need to restrain the animal. Future studies should consider calculating microbiability in rumen, oral and faecal samples from the same animals to directly assess the usefulness of faecal samples comparted to oral and rumen.

### Microbiome functional classification captured methane phenotypic variance

In both trials, microbiability estimates based on functional annotations, including COG and KEGG, were higher than those based on taxonomic classification, even though the differences were not statistically significant as reflected by the large standard errors. Overall, our findings are consistent with previous work reporting higher microbiability with functional classifications (Li et al., 2025). The advantages of functional microbiome representations are well recognised (Xu et al., 2022; Carrascosa et al., 2023), particularly in ecosystems where distinct microbial communities can perform similar metabolic functions, or where community composition is altered by external factors such as temperature and pH without necessarily changing functional capacity. Future studies should examine the optimal bioinformatic and statistical approaches used to both capture microbiome variation, and to use that variation for phenotypic prediction.

In conclusion, this study supports the potential of oral microbiome data as a tool for estimating methane emissions in grazing cattle by showing that the methane emission variation explained using oral microbiome samples is similar to that explained by rumen microbiome samples. Oral microbiome sampling represents a practical, animal-friendly alternative to invasive rumen sampling and to complex methane gas measurement techniques, offering a scalable approach for screening low-methane ruminants in large-scale grazing systems.

## Abbreviations

SOP: standard operating procedures
SF_6_: Sulphur hexafluoride
KEGG: Kyoto Encyclopedia of Genes and Genome database
COG: Clusters of Orthologous Groups
CLR: centred log-ratio

## Acknowledgement

The authors thank Loan To Nguyen and Ziming Chen for their valuable assistance with sample collection and technical support throughout the study. We acknowledge the contributions of colleagues involved in the SF_6_ study that generated the methane dataset and aided with the oral and rumen sampling at the QASP trial. We are also grateful to the staff at Spyglass Research Station,Brian Pastures Research Station for their support with animal handling and on-farm data collection. We gratefully acknowledge The North Australian Pastoral Company and in particular Shane and Sarah Ferriday and team at Goldsborough Station for their support and contribution to this research. This work was supported by Meat & Livestock Australia, whose funding and industry partnership under project P.PSH.2010 made this research possible.

