## Supplementary table 1 for "Short communication: Oral microbiome as a potential proxy for grazing livestock methane emissions"

Supplementary table 1: Summary statistics of sequencing data following quality control, showing the average read N50, quality score, number of reads and total bases across samples after each filtering steps.

| **Bovine oral samples from Brian Pasture, Spyglass and Goldsborough cattle stations** | | | | |
| --- | --- | --- | --- | --- |
| **Step** | **Read N50** | **Mean Qual** | **# Reads (K)** | **Total Bases (Mb)** |
| **Raw** | 1,902.88 | 16.42 | 2,293.16 | 2,138.30 |
| **Chopper** | 2,484.78 | 16.98 | 1,024.52 | 1,737.80 |
| **Host removal** | 3,387.41 | 16.78 | 118.13 | 226.33 |
| **Bovine oral samples from UQ Gatton research trial** | | | | |
| **Step** | **Read N50** | **Mean Qual** | **# Reads (K)** | **Total Bases (Mb)** |
| **Raw** | 1670.22 | 13.18 | 2285.38 | 1851.19 |
| **Host removal** | 1977.41 | 10.24 | 439.86 | 382.85 |
| **Chopper** | 2426.60 | 12.91 | 330.60 | 338.28 |
| **Bovine rumen samples from UQ Gatton research trial** | | | | |
| **Step** | **Read N50** | **Mean Qual** | **# Reads (K)** | **Total Bases (Mb)** |
| **Raw** | 3888.39 | 15.92 | 2241.23 | 3063.40 |
| **Host removal** | 3873.32 | 15.92 | 2218.24 | 3021.46 |
| **Chopper** | 2032.34 | 16.85 | 1736.63 | 1859.18 |
| **Subset** | 2032.32 | 16.85 | 1362.21 | 1497.82 |
